# Anti-integrin αvβ6 autoantibodies are a novel predictive biomarker in ulcerative colitis

**DOI:** 10.1101/2022.11.21.517399

**Authors:** Alexandra E Livanos, Alexandra Dunn, Jeremy Fischer, Ryan C Ungaro, Williams Turpin, Sun-Ho Lee, Shumin Rui, Diane Marie Del Valle, Julia J Jougon, Gustavo Martinez-Delgado, Mark S Riddle, Joseph A Murray, Renee M Laird, Joana Torres, Manasi Agrawal, Jared S Magee, Thierry Dervieux, Sacha Gnjatic, Dean Sheppard, Bruce E Sands, Chad K Porter, Kenneth Croitoru, Francesca Petralia, CCC-GEM Project Research Consortium, OSCCAR Consortium, Jean-Frederic Colombel, Saurabh Mehandru

**Affiliations:** Precision Immunology Institute, Icahn School of Medicine at Mount Sinai, New York, NY, USA; Henry D. Janowitz Division of Gastroenterology, Department of Medicine, Icahn School of Medicine at Mount Sinai, New York, NY, USA; Zane Cohen Centre for Digestive Diseases, Mount Sinai Hospital, Toronto, Ontario, Canada; Division of Gastroenterology & Hepatology, Temerty Faculty of Medicine, University of Toronto, Toronto, Ontario, Canada; Human Immune Monitoring Center, Precision Institute of Immunology, Icahn School of Medicine at Mount Sinai, New York, NY, USA; Hepato-Gastroenterology Department, Claude Huriez Hospital, University of Lille, Lille, France; University of Nevada, Reno - School of Medicine, Reno, Nevada, USA; VA Sierra Nevada Health Care System, Reno, Nevada, USA; Division of Gastroenterology and Hepatology, Mayo Clinic, Rochester, MN, USA; Enteric Diseases Department, Naval Medical Research Center, Silver Spring, MD, USA; Henry M. Jackson Foundation for Military Medicine, Bethesda, MD, USA; Gastroenterology Division, Hospital Beatriz Ângelo, Loures, Portugal; Gastroenterology Division, Hospital da Luz, Lisbon, Portugal; Faculdade de Medicina, Universidade de Lisboa, Portugal; Gastroenterology, Walter Reed National Military Medical Center, Bethesda, USA; Prometheus Laboratories, San Diego, CA, USA; Department of Oncological Sciences, Icahn School of Medicine at Mount Sinai, New York, NY, USA; Division of Pulmonary, Critical Care, Allergy and Sleep, Department of Medicine, University of California, San Francisco, San Francisco, CA, USA; Department of Genetics and Genomic Sciences, Icahn School of Medicine at Mount Sinai, New York, New York, NY, USA

**Author notes:** Corresponding authors Please address correspondence to or. **Author contributions** AL, JFC and SM conceptualized the study, wrote and edited the manuscript. JF developed the anti-αvβ6 ELISA in-house. AL, AD, and JF performed the experiments. All other authors contributed to experimental data and analyses, and along with AL, JFC and SM, they critically reviewed and edited the final version of the manuscript. **Competing interests** SM reports receiving research grants from Genentech and Takeda; receiving payment for lectures from Takeda, Genentech, Morphic; and receiving consulting fees from Takeda, Morphic, Ferring and Arena Pharmaceuticals. JFC reports receiving research grants from AbbVie, Janssen Pharmaceuticals and Takeda; receiving payment for lectures from AbbVie, Amgen, Allergan, Inc. Ferring Pharmaceuticals, Shire, and Takeda; receiving consulting fees from AbbVie, Amgen, Arena Pharmaceuticals, Boehringer Ingelheim, BMS, Celgene Corporation, Eli Lilly, Ferring Pharmaceuticals, Galmed Research, Genentech, Glaxo Smith Kline, Janssen Pharmaceuticals, Kaleido Biosciences, Imedex, Immunic, Iterative Scopes, Merck, Microbia, Novartis, PBM Capital, Pfizer, Protagonist Therapeutics, Sanofi,Takeda, TiGenix, Vifor; and holds stock options in Intestinal Biotech Development RCU has served as an advisory board member or consultant for AbbVie, Bristol Myer Squibb, Janssen, Pfizer, and Takeda; research support from AbbVie, Boehringer Ingelheim, Eli Lily, and Pfizer. TD is employee of Prometheus Laboratories and hold stock options. JAM reports receiving grants from Nexpep/ImmusanT, National Institutes of Health, Immunogenix, Takeda Pharmaceutical, Allakos, Oberkotter, Cour; and consultancy fees from Bionix, Lilly Research Laboratory, Johnson & Johnson, Dr. Schar USA, UCB Biopharma, Celimmune, Intrexon Corporation, Dren Bio, Reistone pharma, Chugai Pharma, Kanyos, Boehringer Ingelheim, Equillium and Torax Medical. SG reports other research funding from Genentech, Boehringer-Ingelheim, EMD Serono, Takeda, and Regeneron. JT received grants from Abbvie and Janssen, payment for lectures from Janssen, Abbvie, and Pfizer, and consulting fees from Janssen, Abbvie, Pfizer, and BMS. DS is a founder of Pliant Therapeutics, receives research funding from AbbVie and is on the Scientific Review Board for Genentech and Amgen. The remaining authors have no competing interests to declare. **Data availability statement** All requests for raw and analyzed data and materials will be promptly reviewed by corresponding author and the study team. We will provide source data files for all the figures. Additional raw data is provided in Supplementary Material. **Code Availability Statement** The R code developed for analyses in this study will be made available.

**Keywords:** Inflammatory bowel disease, Ulcerative Colitis, anti-integrin αvβ6, autoantibodies, biomarkers

## Abstract

**Background and Aims:** Better biomarkers for prediction of ulcerative colitis (UC) development and prognostication are needed. Anti-integrin αvβ6 autoantibodies (anti-αvβ6) have been described in UC patients. Here, we tested for the presence of anti-αvβ6 antibodies in the pre-clinical phase of UC and studied their association with disease-related outcomes after diagnosis.

**Methods:** Anti-αvβ6 were measured in 4 longitudinal serum samples collected from 82 subjects who later developed UC and 82 matched controls from a Department of Defense pre-clinical cohort (PREDICTS). In a distinct, external validation cohort (GEM), we tested 12 pre-UC subjects and 49 matched controls. Further, anti-αvβ6 were measured in 2 incident UC cohorts (COMPASS n=55 and OSCCAR n=104) and associations between anti-αvβ6 and UC-related outcomes were defined using Cox proportional-hazards model.

**Results:** Anti-αvβ6 were significantly higher among individuals who developed UC compared to controls up to 10 years before diagnosis in PREDICTS. The anti-αvβ6 seropositivity was 12.2% 10 years before diagnosis and increased to 52.4% at the time of diagnosis in subjects who developed UC compared with 2.7% in controls across the 4 timepoints. Anti-αvβ6 predicted UC development with an AUC of at least 0.8 up to 10 years before diagnosis. The presence of anti-αvβ6 in pre-clinical UC samples was validated in the GEM cohort. Finally, high anti-αvβ6 was associated with a composite of adverse UC-outcomes including hospitalization, disease extension, colectomy, systemic steroid use and/or escalation to biologic therapy in recently diagnosed UC.

**Conclusion:** Anti-integrin αvβ6 auto-antibodies precede the clinical diagnosis of UC by up to 10 years and are associated with adverse UC-related outcomes.

## Introduction

Inflammatory bowel disease (IBD) including Crohn’s disease (CD) and ulcerative colitis (UC) are chronic inflammatory disorders that primarily affect the gastrointestinal (GI) tract with shared genetic and environmental risk factors^1-3^. Analogous to other immune-mediated diseases, there is an increasing appreciation that IBD may be preceded by sub-clinical immune perturbations^4-6^ and there is a growing interest in biomarkers that can predict the occurrence of IBD.^7^ Additionally, identification of biomarkers to predict disease course is a major unmet need due to significant variability of IBD disease progression^7^. In this context, non-invasive blood-based biomarkers are appealing due to their ease of collection and application in clinical practice. The identification of predictive biomarkers has been more successful in CD^4, 5, 8, 9^, with some being used to risk-stratify patients in clinical trials^10^. In contrast, discovery of predictive biomarkers in UC remains elusive.

Recent studies have highlighted a novel autoantibody against integrin αvβ6 in the serum of patients diagnosed with UC^11-13^. In a study from Japan, anti-integrin αvβ6 (anti αvβ6) autoantibodies had a sensitivity of 92.0% and a specificity of 94.8% for diagnosing UC in adult patients as compared to non-IBD subjects^11^. These results were further confirmed in a Swedish cohort and extended to a Japanese pediatric population^12, 13^. Integrin αvβ6 is an epithelium-associated heterodimer that interacts with the extracellular matrix^14^ and enables the activation of latent TGF-β, which is thought to be its primary role *in vivo*^15, 16^. Accordingly, critical homeostatic roles such as maintenance of epithelial barrier integrity and suppression of epithelial inflammation have been ascribed to integrin αvβ6^17-19^.

Recognizing that loss of epithelial barrier integrity is a major and perhaps early feature of disease pathogenesis^20^, we hypothesized that onset of UC may be preceded by the appearance of anti-αvβ6 autoantibodies and thus, may serve as a pre-clinical biomarker. Furthermore, due to the important physiological role of integrin αvβ6, we explored the performance of anti-αvβ6 autoantibodies to predict adverse clinical outcomes in recently diagnosed UC.

Our study leverages two unique pre-clinical cohorts of UC. In our primary pre-clinical cohort, PRoteomic Evaluation and Discovery in an IBD Cohort of Tri-service Subjects (PREDICTS), longitudinal serum samples were obtained up to 10 years prior to disease diagnosis. The Crohn’s and Colitis Canada Genetics Environment Microbial (GEM) Project, served as an independent external validation pre-clinical cohort. Finally, we also studied two well-characterized incident IBD cohorts to further understand the association between anti-αvβ6 and UC-related disease outcomes.

## Materials and Methods

### Clinical Cohorts

#### PREDICTS cohort, primary pre-clinical cohort

We studied longitudinal serum samples from 82 individuals who eventually developed ulcerative colitis (UC) and 82 healthy controls (HC) matched by age, gender and race from the PREDICTS cohort which was created in collaboration with the Department of Defense Serum Repository (DoDSR)^21^. This study was approved by the institutional review board of the Naval Medical Research Center (NMRC.2014.0019), Silver Spring, MD in compliance with all Federal regulations governing the protection of human volunteers and this research was performed under a Cooperative Research and Development Agreement (NMR 17-10209). Incident cases of UC between 1998 and 2013 within the Defense Medical Surveillance System (DMSS) were identified by having 2 or more medical encounters with the ICD-9 code for UC [556 (including all subgroup codes) and linked to the DoDSR]. For each subject, 4 serum samples were retrieved: one from the time of diagnosis (± 1 year) (Sample A) and 3 other samples preceding diagnosis (Sample B, C and D). HC were required to have no medical encounter with evidence of IBD, rheumatoid arthritis, celiac disease, or colon cancer (based on ICD-9 codes) and available serum at the time of Sample A for UC subjects (±1 year). Controls were matched based on age, gender and race. Sample B was at a median of -2.01 years before diagnosis (Sample A) for UC and -2.03 (before Sample A) years for HC, Sample C was at a median of -4.06 for UC and -3.99 years for HC, and Sample D, the earliest available sample, was at a median of -10.03 for UC and -10.52 for HC. Measurement of anti-integrin αvβ6 IgG autoantibodies and total IgG was performed with blinding to diagnosis and sample timepoint. From Sample D, 1 serum sample from a HC subject was excluded due to discrepancy in serum ID identified after unblinding. ICD-9 codes considering the code representing the highest disease extent were used to determine the disease extent of the UC patients according to the Montreal classification ^4, 9, 22^. Specifically, the extent of disease was defined by 3 subgroups: E1 proctitis only (disease limited to the rectum), E2 left sided colitis (disease distal to the splenic flexure), and E3 extensive colitis (disease beyond the splenic flexure)^22^.

#### GEM cohort, validation pre-clinical cohort

We evaluated 61 sera samples that were collected as part of the prospective CCC-GEM Project. As described elsewhere^5, 23^, this is a prospective study which recruited asymptomatic first-degree relatives (FDR) of patients with CD between 2008 and 2017. Subjects were between the age of 6 and 35 at the time of recruitment. The study was approved by the Mount Sinai Hospital Research Ethics Board (Toronto-Managing Center) and the local recruitment centers. All subjects were contacted every 6 months via a phone call and if a subject reported being diagnosed with UC, this was confirmed by their treating physician based on clinical, endoscopic, radiographic, and/or histologic reports. The present study represents a nested case-control within the CCC-GEM, including 12 individuals who developed UC (cases) matched to at least 4 controls per subject, by age, gender, geographic location (by postal and country codes), and follow-up duration. As with the PREDICTS cohort samples, all samples from the GEM cohort were blinded during sample testing.

#### COMPASS cohort

The Comprehensive Care for the Recently Diagnosed IBD Patients (COMPASS) cohort is a prospective cohort of recently diagnosed patients with IBD who are enrolled in a registry at The Mount Sinai Hospital (MSH, New York, NY) and approved by the institutional review board (STUDY-17-01304). All patients are enrolled within 18 months of diagnosis and disease diagnosis was confirmed based on standard criteria^24^. Baseline demographic and clinical variables were obtained by patient questionnaires and standardized data abstraction from the medical record including age, sex, race/ethnicity, baseline medications, laboratory data including c-reactive protein (CRP) and erythrocyte sedimentation rate (ESR), and disease extent (following the Montreal classification^22^). Outcomes were determined by chart review by a gastroenterologist to determine if the patient had IBD-related hospitalization, extension of disease (defined as E1 disease extending to either E2 or E3, or E2 disease extending to E3 disease), IBD-related surgery, systemic steroid use (defined as oral prednisone or intravenous steroid formulations) or required biologic therapy. Serum samples from the time of enrollment were available on 55 UC patients which were used in this study. Non-IBD control subjects were defined as subjects without an IBD diagnosis who were seen at MSH GI clinic or endoscopy and had serum or plasma samples available for testing.

#### OSCCAR cohort

The Ocean State Crohn’s and Colitis Area Registry (OSCCAR) is a community-based prospective inception cohort established in the state of Rhode Island (Sands Med Health Rhode Island) as previously described^25, 26^. Enrollment was between January 2008 and 2018. Patients were included if they had a confirmed new diagnosis of IBD by endoscopic, pathologic, or radiographic findings according to the criteria of the NIDDK IBD Genetics Consortium^2^ and were a resident of Rhode Island at the time of diagnosis. Patients were excluded if they were previously diagnosed with IBD or were unwilling to provide informed consent. Baseline and longitudinal data including medication prescriptions, IBD-related hospitalizations, IBD-related surgeries, and endoscopic disease extent (based on the Montreal classification^22^) were collected by annual structured interview, self-completion of validated questionnaires, and standardized central data abstraction from the medical record. Additionally, select laboratory data was available including CRP and ESR as well as perinuclear anti-neutrophil cytoplasmic antibodies (pANCA) at baseline. pANCA testing was performed at Prometheus Laboratories (San Diego, CA). In this cohort, there were 151 UC subjects. Two patients were eliminated as they did not have follow up data and 1 additional subject was removed as diagnosis > 6 months from enrollment. Therefore, analysis was performed on the remaining 148 samples. Of these 148 subjects, 143 had pANCA data to analyze and 104 had serum available for measurement of anti-integrin αvβ6 IgG autoantibodies.

### ELISA for measurement of total IgG and anti-integrin αvβ6 IgG autoantibodies

Total serum IgG was measured using Mabtech human IgG ELISA kit following the manufacturer’s protocol. For detection of IgG against integrin αvβ6, all serum samples were diluted to a concentration of 10 μg/ml IgG for subsequent testing for anti-integrin αvβ6 IgG as detailed below. MaxiSorp immuno microtiter plates (Thermo Fisher Scientific) were coated with 100 μl per well of 1.5 μg/ml human recombinant integrin αvβ6 (R&D Systems) diluted in coating buffer from the ELISA Starter Accessory Kit (Bethyl Laboratories) and incubated at room temperature for 60 minutes. Next, 1 mM MgCl2 and CaCl_2_ were added to the incubation buffers to stabilize the αvβ6 integrin heterodimer as described by Kuwada et al^11^. The plates were subsequently washed with ELISA washing buffer and blocked with blocking buffer for 30 minutes at room temperature (all buffers purchased from Bethyl Laboratories). The plates were then washed and incubated with 100 μl per well of samples prepared as described above and with mouse anti-human αvβ6 (Millipore, MAB2077Z) from 312.5 ng/ml to 1.22 ng/ml to produce the standard curve for 60 min at room temperature. Plates were then washed and incubated with secondary antibodies. For the human sera samples, anti-human IgG secondary conjugated to horseradish peroxidase (HRP) (Invitrogen, 31410) diluted 1:4000 was used, and for the standards, anti-mouse IgG secondary conjugated to HRP (Invitrogen, 62-6520) diluted 1:2000 was used. Plates were washed and then developed by incubating with 100 μl per well of 3,3′,5,5′-tetramethylbenzidine for 4 minutes, at which time the reaction was stopped with 100 μl per well of 0.18 M H_2_SO_4_ and immediately read on the POLARstar Omega plate reader (BMG LABTECH).

In a subset of patients with established UC (n=6) and HC (n=2), plasma samples were assayed for anti-integrin αvβ6 IgG at a 1:200 starting dilution and 7 additional threefold serial dilutions from which the area under the curve (AUC) was calculated. In addition, we measured anti-αvβ6 IgG as described above with samples diluted to a concentration of 10 μg/ml IgG (IgG normalized measurement). The IgG normalized measurement strongly correlated with the AUC from the dilution curves (Pearson’s correlation r=0.97, p<0.0001) (**Supplementary Figure 1 A** and **B**)

### Statistical Analysis

Descriptive statistics were performed in GraphPad Prism (version 9.3.0). The anti-αvβ6 IgG was treated as a continuous variable and binary variable. As a binary variable the cut-off for positivity was defined as greater than the mean HC plus 3 standard deviations (SD)^11^. For the univariate analyses of continuous variables, we used the Mann-Whitney test (for 2 group analyses) or Kruskal Wallis with Dunn’s multiple comparisons test (for >2 group analyses). For categorical comparisons, the Fisher’s exact or chi-square test were used. For comparing the anti-αvβ6 IgG OD450 between pre-UC and matched HC (or asymptomatic) subjects in the PREDICTS cohort and the GEM cohort we performed conditional logistic regressions using the R function *clogit* from the survival package^27^. Linear regression using g*lm* function in r was used to model anti-αvβ6 as a function of the disease extent in the PREDICTS cohort^28^. Spearman’s correlations were used to determine strength and direction of the associations between two variables.

In order to assess the predictive performance of anti-αvβ6 IgG in the PREDICTS cohort, 10-fold cross validation was performed. For each fold, the model was trained on the remaining 9 set of samples and the predictive performance of the estimated model was applied to predict the outcome of samples in the validation set. For this analysis, UC status was modeled via logistic regression as a function of anti-αvβ6. Predictive performance was evaluated based on Receiver Operating Characteristic Curves (ROC) and AUC. Logistic regressions were estimated using the R function *glm*^28^. 95% confidence interval (CI) of AUC were evaluated based on 10,000 bootstrap iterations using function ci.auc from the pROC package^29^ available in R.

For the COMPASS and OSCCAR cohort analyses, multivariable regressions was performed using the R function *glm*^28^ and ROC curves created in R using the pROC package^29^ and graphed in GraphPad Prism. 95% CI of AUC were evaluated as described above with 10,000 bootstrap iterations. Cox proportional-hazards models were used to define associations between anti-αvβ6 and UC-related outcomes and visualized using Kaplan-Meier curves. For these analyses we used a composite of disease-related adverse outcomes including the need for biologic therapy, disease extension, systemic steroid use, UC-hospitalization and / or surgery. In the COMPASS model, we used age and extensive disease (E3) at baseline as covariates. In the OSCCAR model, we included the clinical risk factors for complicated disease course including age <40 years old at diagnosis, extensive disease (E3), ESR / CRP elevation, systemic steroid use, focal ulcers and history of UC-related hospitalization^30^.

## Results

### *Anti-*αvβ6 *autoantibodies are detected up to 10 years prior to the diagnosis of UC*

We hypothesized that anti-αvβ6 autoantibodies would predate UC diagnosis and analyzed longitudinal samples predating UC diagnosis by up to 10 years in 82 subjects who developed UC matched by age, gender and race to 82 subjects that did not develop IBD (HC) (**Figure 1A** and **Table 1**). Anti-αvβ6 levels were significantly higher in sera from patients who developed UC compared to the controls at all timepoints tested (p <0.001 Sample A-C, p=0.0015 Sample D, **Supplementary Table 1**) (**Figures 1B** and **C**). During the pre-clinical phase (Sample D to A), anti-αvβ6 seropositive subjects increased from 12.2% (Sample D) to 20.7% (Sample C) to 30.5% (Sample B) to 52.4% (Sample A) of subjects who developed UC (Chi-square test for trend p<0.0001), compared with a mean of 2.7% in HC group across the 4 timepoints (**Figure 1C**).

**Figure 1:**
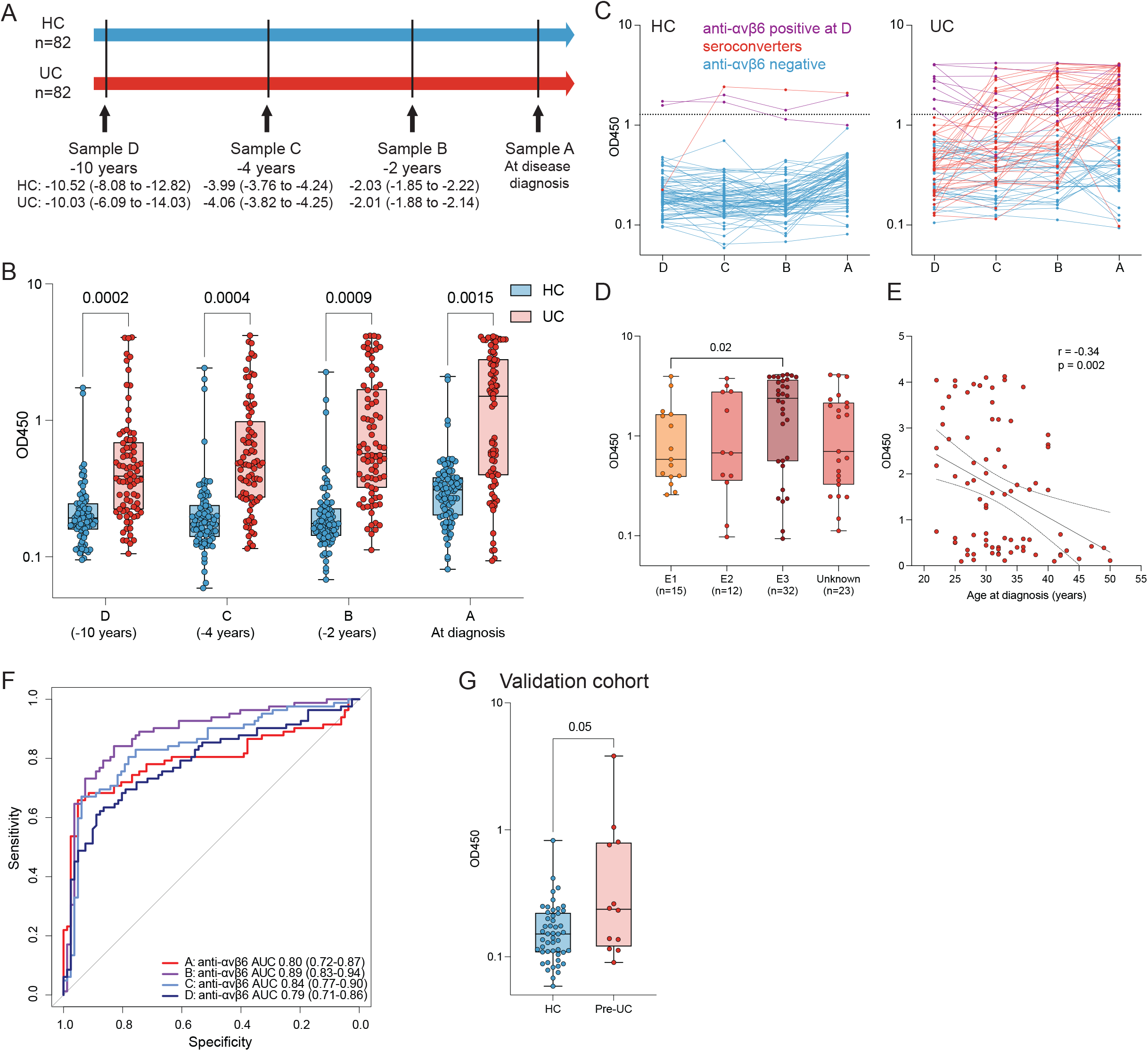
Anti-αvβ6 autoantibodies in pre-clinical ulcerative colitis subjects. **(A)** Timeline of sample collection in the PREDICTS cohort from 82 subjects who develop ulcerative colitis (UC) and 82 matched controls (HC). Sample A was collected at the time of diagnosis for UC subjects and Sample B, C and D were collected prior to diagnosis. The median time in years as well as interquartile range (IQR) for Sample A, B, C and D are detailed for both UC and HC. **(B)** Anti-αvβ6 autoantibody absorbance values (OD450) determined by ELISA in UC (n=82, shown in red) and HC (n=82, shown in blue) sera obtained prior to diagnosis (Sample B, C and D) and at the time of diagnosis (Sample A) from the PREDICTS cohort. In Sample D, 1 HC sample was excluded due to identification issue. For all boxplots, the box represents the IQR, the center line represents the median and the whiskers show the minimum to maximum value. Statistical significance determined by conditional logistic regressions at each timepoint comparing subjects that developed UC with matched controls. **(C)** Dynamics of anti-αvβ6 in each HC or UC subject across the 4 longitudinal samples in the PREDICTS cohort. Purple lines represent samples that are positive for anti-αvβ6 in Sample D, red lines indicate samples that seroconvert between Sample D and A and blue lines indicate subjects that are negative at all sample timepoints. **(D)** Anti-αvβ6 autoantibody levels as a function of UC disease extent as defined by Montreal classification (E1: proctitis only (n=20), E2: Left-sided disease (n=15), E3: extensive disease (n=40), unknown (n=25)), and **(E)** by age at diagnosis shown as scatterplot in the PREDICTS cohort. Statistical significance determined by linear regression and Spearman’s correlation, respectively. **(F)** Predictive performance of anti-αvβ6 autoantibodies based on 10-fold cross validation and the 95% confidence interval (CI) of the area under the curve (AUC) based on 10000 bootstrap iterations in the PREDICTS cohort. Sample A (time of diagnosis) is in red, Sample B (−2 years) is in purple, Sample C (−4 years) is in light blue, and Sample D (−10 years) is in dark blue. **(G)** Anti-αvβ6 autoantibody absorbance values (OD450) determined by ELISA in 12 pre-UC subjects and 49 matched control subjects from the GEM cohort. Statistical significance determined by conditional logistic regression comparing pre-UC subjects with matched control subjects.

**Table 1:** Demographic and Clinical Characteristics of PREDICTS Cohort.

|  | Healthy controls | Ulcerative colitis |
| --- | --- | --- |
| Number of individuals | 82 | 82 |
| Number of samples | 327* | 328 |
| Age in years at diagnosis or sample A, mean ± SD | 32.6 ± 5.6 | 32.0 ± 6.4 |
| Male (n) / Female (n) | 79 / 3 | 79 / 3 |
| Race, n |  |  |
| White | 62 | 62 |
| Black | 18 | 18 |
| Other | 2 | 2 |
| Disease extent at diagnosis, n |  |  |
| E1, proctitis | NA | 15 |
| E2, Left-sided colitis | NA | 12 |
| E3, extensive | NA | 32 |
| unknown | NA | 23 |

\*Sample D from 1 HC subject excluded due to ID discrepancy
SD = Standard deviation

Next, we examined disease-related factors that were associated with anti-αvβ6 (in Sample A, at diagnosis), and determined that anti-αvβ6 were associated with extensive (E3) disease (odds ratio (OR) 2.76, 95% CI, 1.21-6.33, **Figure 1D**) and inversely associated with age of diagnosis (Spearman’s correlation r=-0.34, p=0.002, **Figure 1E**).

After identifying the presence of anti-αvβ6 in UC samples prior to diagnosis, we sought to determine the performance of this autoantibody as a predictive biomarker. The predictive performance of anti-αvβ6 was assessed via ROC curves and AUC based on 10-fold cross validation. The AUC was 0.80 at the time of diagnosis (Sample A). Notably, the AUC remained high in all three pre-diagnostic samples with AUC of 0.89, 0.84 and 0.79 at Sample B, C and D, respectively (**Figure 1F** and **Table 2**). Further, the lower bound of the 95% CI of AUCs calculated via bootstrapping were above 0.71 for samples A, B, C and D; confirming the excellent predictive performance of anti-αvβ6 in all pre-diagnostic groups of samples (**Figure 1F** and **Table 2**).

**Table 2:**
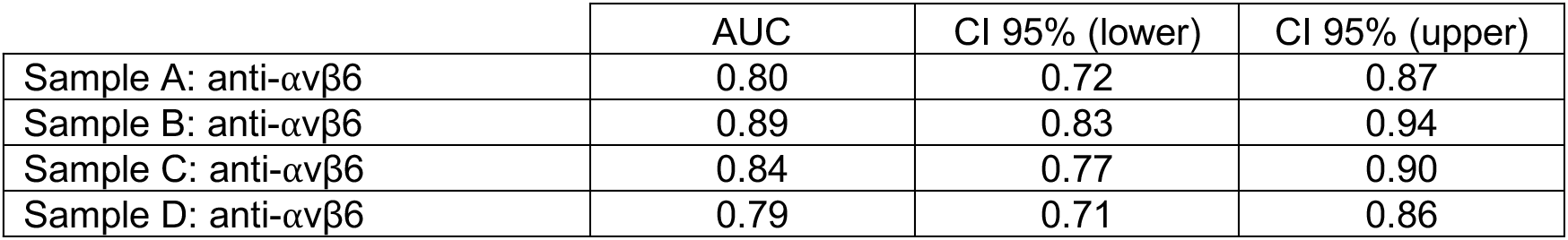
Predictive model for UC development in PREDICTS samples. Area under the receiving characteristic curves (AUC) and corresponding 95% confidence intervals (CI) based on 10-fold cross validation.

To confirm our findings in a second independent cohort, we tested sera collected as part of the CCC-GEM Project. In this cohort of >5000 FDRs of CD patients, 12 subjects had developed UC since recruitment (median time from recruitment to diagnosis was 4.2 years (range 0.4 to 8.5)). We tested these pre-UC samples and compared them with 49 matched subjects who remained asymptomatic (HC) since recruitment (**Table 3**). The pre-UC samples had higher anti-αvβ6 autoantibodies compared with controls (conditional logistic regression p=0.05) (**Figure 1G**). In the pre-UC group, 4 out of the 12 subjects (33%) were positive for the anti-αvβ6 (defined as mean + 3 SD of asymptomatic subjects from the GEM cohort) while 1 of the 49 of controls (2%) were anti-αvβ6 positive (Fisher’s exact p=0.004).

**Table 3:** Demographic and Clinical Characteristics of GEM cohort.

|  | Subjects that remained asymptomatic (HC) | Subjects that developed UC (pre-UC) |
| --- | --- | --- |
| Number of individuals | 49 | 12 |
| Age in years at recruitment, mean ± SD | 17.7 ± 7.1 | 17.3 ± 7.5 |
| Male (n) / Female (n) | 24 / 25 | 6 / 6 |
| Country recruitment, n |  |  |
| Canada | 40 | 10 |
| Israel | 5 | 1 |
| USA | 4 | 1 |
| Relation to proband, n |  |  |
| Offspring | 23 | 1 |
| Sibling | 26 | 11 |
| Time from recruitment to diagnosis in years, mean (range) | NA | 4.2 (0.4-8.5) |

SD = Standard deviation

### *Anti-*αvβ6 *autoantibodies are associated with adverse disease related outcomes*

Due to the physiological roles ascribed to αvβ6 in mucosal homeostasis, we hypothesized that anti-αvβ6 autoantibodies would be associated with adverse clinical outcomes. To test this hypothesis, we studied anti-αvβ6 in two inception cohorts of patients with UC, COMPASS (n=55) and OSCCAR (n=104), and compared them with non-IBD controls (non-IBD) (n=54), **Table 4**. With the exception of the UC patients being younger than controls (p<0.0001 COMPASS, p=0.0011 OSCCAR), cases and controls shared similar characteristics. Ninety-one percent of COMPASS patients and 96% of OSCCAR patients were naïve to any biologic therapies.

**Table 4:** Demographics of two ulcerative colitis inception cohorts and non-IBD controls.

|  | Non-IBD controls | COMPASS UC | OSCCAR UC |
| --- | --- | --- | --- |
| Number of individuals | 54 | 55 | 104 |
| Number of samples | 54 | 55 | 104 |
| Age in years at diagnosis ± SD | 46.3 ± 15.9 | 31.3 ± 12.4 | 36.8 ± 20.1 |
| Male (n) / Female (n) | 31 / 23 | 28 / 27 | 42 / 62 |
| Race, n |  |  |  |
| White | 19 | 35 | 92 |
| Black | 14 | 4 | 4 |
| Asian | 3 | 5 | 0 |
| Other | 18 | 11 | 8 |
| Disease extent at diagnosis, n |  |  |  |
| E1, proctitis | NA | 7 | 27 |
| E2, Left-sided colitis | NA | 24 | 66 |
| E3, extensive | NA | 24 | 44 |
| Mayo Endoscopic score, n |  |  |  |
| 0 | NA | 5 | NA |
| 1 | NA | 15 | NA |
| 2 | NA | 16 | NA |
| 3 | NA | 14 | NA |
| Unknown | NA | 5 | 104 |
| Biologic naïve n (%)<br>(at enrollment/blood draw) | 54 (100%) | 50 (90.9%) | 100 (96.2%) |

NA = not applicable or not available

Anti-αvβ6 levels were significantly higher in both the COMPASS and OSCCAR UC patients compared to non-IBD controls (p<0.0001) (**Figure 2A**) consistent with published data.^11, 13^ ROC analysis revealed an AUC of 0.99 (95% CI 0.97-1.00 for COMPASS) and AUC 0.96 (95% CI 0.93-0.98) for OSCCAR (**Figure 2B**). With a cut-off of 0.99 (mean non-IBD + 3SD), the sensitivity of anti-αvβ6 for distinguishing UC from non-IBD controls was 85.5% and 70.2% for the COMPASS cohort and for the OSCCAR cohort respectively with specificity of 98.1%. Anti-αvβ6 remained significantly associated with UC diagnosis in a multivariable model including age, sex, and race (OR 64.05, 95% CI 7.41-553.74, p=0.0002 for COMPASS and OR 156.29, 95% CI 12.53-1949.04, p=0.0001 for OSCCAR) (**Supplementary Table 2** and **3**).

**Figure 2:**
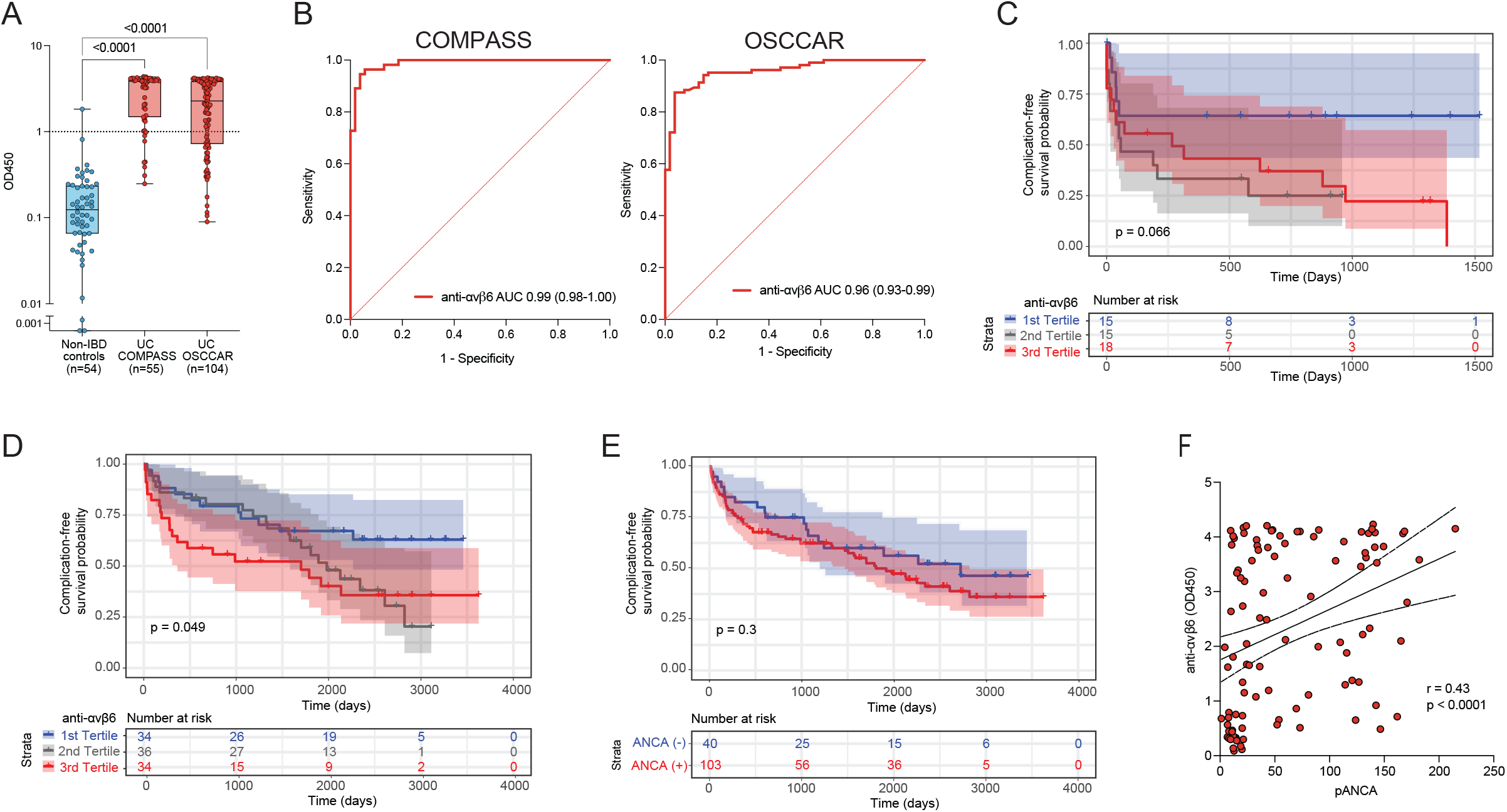
Anti-αvβ6 autoantibodies in patients with newly diagnosed ulcerative colitis and their association with adverse disease-related outcomes. **(A)** Anti-αvβ6 autoantibody absorbance values (OD450) determined by ELISA in non-IBD controls (n=54, shown in blue), in COMPASS UC patients (n=55, shown in red) and in OSCCAR UC patients (n=104, shown in red). **(B)** Receiver operator curve analysis of anti-αvβ6 autoantibodies in both COMPASS UC patients (left) and OSCCAR UC patients (right) compared with the non-IBD controls. **(C)** and **(D)** Kaplan-Meier curve for composite outcome of IBD hospitalization, proximal disease extension, need for surgery, systemic steroid use and / or requiring new biologic therapy in the COMPASS cohort (**C**) and OSCCAR cohort (**D**) stratified by anti-αvβ6 titer tertiles. The blue line represents the 1^st^ tertile (lowest), the grey line the 2^nd^ tertile and the red line the 3^rd^ (highest) tertile. **(E)** Kaplan-Meier curve for the same composite outcomes in the OSCCAR cohort stratified by the presence or absence of pANCA. **(F)** Spearman’s correlation between pANCA and anti-αvβ6 titers.

Next, we examined if anti-αvβ6 autoantibodies were associated with any UC-related adverse outcomes representing more complicated disease defined as a composite of need for biologic therapy, disease extension, systemic steroid use, IBD-related hospitalization, and / or surgery. In the COMPASS cohort, we found that anti-αvβ6 autoantibodies were significantly associated with the composite of the above disease-related outcomes (Hazard ratio (HR) 1.39, 95% CI 1.03-1.89, with inclusion of disease extent adjusted HR 1.35, 95% CI 0.99-1.85) (**Figure 2C, Supplementary Table 4)**. In the OSCCAR cohort, we confirmed these findings and again found that subjects with higher autoantibodies were more likely to have a more complicated course as defined by the same composite outcome even when correcting for baseline clinical risk factors for complications including age <40 years old at diagnosis, extensive disease (E3), elevated CRP or ESR, use of systemic steroids or hospitalization at diagnosis, and ulcers on baseline endoscopy^30^ (adjusted HR 1.24, 95% CI 1.01-1.53) (**Figure 2D, Supplementary Table 5**). In contrast, pANCA, an established UC-associated biomarker^31, 32^, was not associated with disease outcomes (HR 0.75, 95% CI 0.36-1.54) (**Figure 2E, Supplementary Table 6**); however, we did note a moderate positive correlation between anti-αvβ6 and pANCA (r=0.43, p<0.0001) (**Figure 2F**).

## Discussion

Here, we identify anti-αvβ6 autoantibodies as a new pre-clinical biomarker that precedes the diagnosis of UC by up to 10 years. Further, we provide evidence in support of anti-αvβ6 autoantibodies as a prognostic biomarker associated with adverse UC-related outcomes in two well-characterized incident IBD cohorts of recently diagnosed patients.

Prior work had shown that anti-αvβ6 autoantibodies were present in 92% of patients with established UC and had a high sensitivity and specificity for the diagnosis of UC^11^. The present study reports that anti-αvβ6 autoantibodies are associated with the pre-clinical phase of UC. Importantly, these data were replicated in an independent external validation cohort.

The predictive performance of anti-αvβ6 autoantibodies as assessed via AUROC with 10-fold cross validation was at least 0.8 at all timepoints and remained predictive even up to 10 years before diagnosis. Furthermore, the number of seropositive patients significantly increased prior to the development of clinically overt disease. The increasing prevalence of anti-αvβ6 autoantibodies with time is in contrast to other autoantibodies such as ASCA and pANCA which remain relatively stable prior to diagnosis^4^. The predictive performance of anti-αvβ6 is superior to that of pANCA, an established UC-associated diagnostic biomarker^4^. Specifically, pANCA was found to be positive in a small subset of pre-clinical UC subjects (2/8, 25% in Israeli et al)^33^ but had poor predictive performance with AUCs in the range of 0.59-0.61^4^. The combination of anti-microbial antibodies and pANCA also had a low predictive performance for UC, with AUCs ranging from 0.57 at 5 years to 0.61 at 1 year before diagnosis in a prior study using samples derived from the PREDICTS cohort. In the same study, 1129 proteins were measured using the SomaLogic platform and predictive performances using select disease-associated markers also were suboptimal with AUCs ranging from 0.49 5 years before diagnosis to 0.68 at diagnosis^4^. In the EPIC (European Prospective Investigation into Cancer and Nutrition) study which included a single pre-diagnostic sample from 54 subjects who developed UC a mean of 4.4 years after collection, a combination of anti-microbial antibodies (including pANCA, anti-CBir1, anti-OmpC and ASCA IgA) had an AUC <0.70^34^. Additionally, in the Nurses’ Health Study cohort, subjects with the highest quintile of serum interleukin-6 and high-sensitivity CRP were more likely to develop UC with an OR 3.43 (95% CI 1.44-8.15) and 1.79 (95% CI 0.80-3.99), respectively; however, these associations were only present at the highest quintiles^35^. Finally, in a population cohort from Sweden, six proteins (MMP10, CXCL9, CCL11, SLAMF1, CXCL11, and MCP-1) were increased in patients who later developed UC compared to those that remained healthy, with a high predictive performance (AUC 0.92); however, the predictive performance of these proteins was assessed in an incident cohort rather than in a pre-clinical cohort^6^. Thus, to date, anti-αvβ6 autoantibodies have at least as high (or higher) predictive performance as all existing UC-associated biomarkers and may accurately identify patients at risk for developing UC. The presence of anti-αvβ6 autoantibodies before UC diagnosis further suggests that a pre-clinical phase, potentially amenable to therapeutic intervention may indeed predate UC diagnosis.

Next, we examined anti-αvβ6 autoantibodies in two well-characterized cohorts of recently diagnosed IBD, COMPASS and OSCCAR, and found a significant association of this autoantibody with adverse UC-related outcomes that included disease extension, escalation to biologic therapy, need for systemic steroids, and UC-related surgery, and / or hospitalization. Consistent with prior data^36, 37^, pANCA was not associated with these adverse clinical outcomes. Another potentially promising non-invasive biomarker is a panel of serum proteins which associated with the need for treatment escalation in both UC and CD^38^. This serum protein panel is now being investigated in the Nordic IBD treatment strategy trial (NORDTREAT) which includes patients with UC as well as CD. Notably, the five-protein model with the highest predictive accuracy included ITGAV and EpCAM^38^. Interestingly, EpCAM is an epithelial cell-associated adhesion molecule ^39^ and ITGAV is the integrin subunit alpha V which is part of the alphaVbeta6 (αvβ6) integrin heterodimer that is targeted by anti-αvβ6 autoantibodies^11, 14^. ITGAV has also been identified as an IBD risk allele in GWAS studies^40^. Altogether, following confirmation in larger data sets, anti-αvβ6 can potentially serve to risk-stratify UC patients to identify those who may benefit from earlier introduction of targeted immunosuppressive therapy and can be studied in a ‘biomarker-stratified’ trial similar to the PROFILE trial in CD^10^.

Considering the strong predictive value of anti-αvβ6 autoantibodies for UC, understanding the pathophysiological role of these autoantibodies could bring light some of the early pathophysiological events associated with this disease. For example, we have recently identified colonic plasma cells specific to integrin αvβ6 in a patient with UC^41^, providing evidence that these autoantibodies originate from intestinal plasma cells. αvβ6 integrin expression is low in the adult intestinal epithelium but dramatically increases in setting of epithelial injury which possibly leads to increased exposure of this autoantigen^16, 42, 43^. Additionally, *in vitro* studies from Kuwada et al demonstrate that anti-αvβ6 autoantibodies inhibit binding of integrin αvβ6 to fibronectin^11^, and therefore, may disrupt homeostatic epithelial-stromal interactions, culminating in impaired barrier integrity^19^. Other potential mechanisms of action of anti-αvβ6 may be mediated through blockade of the activation of TGF-β resulting in increased inflammation^15^ and antibody-dependent cellular cytotoxicity, an important mechanism of action of autoantibodies in other autoimmune conditions such as vitiligo^44^.

Our study has several strengths including the detection of anti-αvβ6 in two independent pre-clinical UC cohorts. In our primary pre-clinical cohort, PREDICTS, multiple longitudinal pre-diagnostic samples allowed us to study the evolution of anti-αvβ6 autoantibodies over time. Additionally, we have provided evidence of anti-αvβ6 autoantibodies as a potential prognostic biomarker using two independent incident cohorts. It is possible that subjects had unrecognized symptoms of UC prior to the clinical diagnosis in the PREDICTS cohort. However, we believe this is limited given the routine medical evaluations and physical rigor required of active component military populations. Furthermore, this potential limitation does not apply to samples collected up to 10 years before diagnosis and subjects in our pre-clinical validation cohort (GEM) which were required to be asymptomatic at enrollment. Finally, we were unable to perform functional studies due to limited sample availability.

In conclusion, anti-αvβ6 autoantibodies are a novel biomarker associated with the pre-clinical phase of UC and are a prognostic biomarker associated with development of complicated disease in recently diagnosed patients. These findings further highlight potential key early events in the development of UC providing opportunities for diagnostic and therapeutic interventions.

## Acknowledgements

We acknowledge the contributions of the members of the CCC-GEM Project Research Consortium as well as the OSCCAR Consortium including Jason Shapiro, Samir Shah and Neal S. Leleiko. The CCC-GEM Project Research Consortium is composed of: Maria Abreu, Paul Beck, Charles Bernstein, Kenneth Croitoru, Leo Dieleman, Brian Feagan, Anne Griffiths, David Guttman, Kevan Jacobson, Gilaad Kaplan, Denis O. Krause* (*deceased), Karen Madsen, John Marshall, Paul Moayyedi, Mark Ropeleski, Ernest Seidman*, Mark Silverberg, Scott Snapper, Andy Stadnyk, Hillary Steinhart, Michael Surette, Dan Turner, Thomas Walters, Bruce Vallance, Guy Aumais, Alain Bitton, Maria Cino, Jeff Critch, Lee Denson, Colette Deslandres, Wael El-Matary, Hans Herfarth, Peter Higgins, Hien Huynh, Jeff Hyams, David Mack, Jerry McGrath, Anthony Otley, and Remo Panancionne. The CCC-GEM Project recruitment site directors include Maria Abreu, Guy Aumais, Robert Baldassano, Charles Bernstein, Maria Cino, Lee Denson, Colette Deslandres, Wael El-Matary, Anne M. Griffiths, Charlotte Hedin, Hans Herfarth, Peter Higgins, Seamus Hussey, Hien Hyams, Kevan Jacobson, David Keljo, David Kevans, Charlie Lees, David Mack, John Marshall, Jerry McGrath, Sanjay Murthy, Anthony Otley, Remo Panaccione, Nimisha Parekh, Sophie Plamondon, Graham Radford-Smith, Mark Ropeleski, Joel Rosh, David Rubin, Michael Schultz, Ernest Seidman*, Corey Siegel, Scott Snapper, Hillary Steinhart, and Dan Turner.

## Department of Defense Statement

### Disclaimer

The views expressed in this article are those of the authors and do not necessarily reflect the official policy or position of the Department of the Navy, Department of the Army, Department of Defense, nor the U.S. Government. This is a US Government work. There are no restrictions on its use. There were no financial conflicts of interests among any of the authors.

### Copyright Statement

One of the authors (CKP) is an employee of the U.S. Government. This work was prepared as part of their official duties. Title 17 U.S.C. §105 provides that “Copyright protection under this title is not available for any work of the United States Government.” Title 17 U.S.C. §101 defines a U.S. Government work as a work prepared by a military service member or employee of the U.S. Government as part of that person’s official duties.

## References

1. Bonen DK, Cho JH. The genetics of inflammatory bowel disease. Gastroenterology 2003;124:521–36.

2. NIDDK IBD Genetics Consortium Phenotype Operating Manual. 2006 May 10. http://info.med.yale.edu/intmed/ibdgc/resources/docs/Phenotyping_Manual_5-10-2006.pdf.

3. Ungaro R, Mehandru S, Allen PB, et al. Ulcerative colitis. Lancet 2017;389:1756–1770.

4. Torres J, Petralia F, Sato T, et al. Serum Biomarkers Identify Patients Who Will Develop Inflammatory Bowel Diseases Up to 5 Years Before Diagnosis. Gastroenterology 2020;159:96–104.

5. Lee SH, Turpin W, Espin-Garcia O, et al. Anti-Microbial Antibody Response is Associated With Future Onset of Crohn’s Disease Independent of Biomarkers of Altered Gut Barrier Function, Subclinical Inflammation, and Genetic Risk. Gastroenterology 2021;161:1540–1551.

6. Bergemalm D, Andersson E, Hultdin J, et al. Systemic Inflammation in Preclinical Ulcerative Colitis. Gastroenterology 2021;161:1526–1539 e9.

7. Verstockt B, Parkes M, Lee JC. How Do We Predict a Patient’s Disease Course and Whether They Will Respond to Specific Treatments? Gastroenterology 2022;162:1383–1395.

8. Lee JC, Lyons PA, McKinney EF, et al. Gene expression profiling of CD8+ T cells predicts prognosis in patients with Crohn disease and ulcerative colitis. J Clin Invest 2011;121:4170–9.

9. Choung RS, Princen F, Stockfisch TP, et al. Serologic microbial associated markers can predict Crohn’s disease behaviour years before disease diagnosis. Aliment Pharmacol Ther 2016;43:1300–10.

10. Parkes M, Noor NM, Dowling F, et al. PRedicting Outcomes For Crohn’s dIsease using a moLecular biomarkEr (PROFILE): protocol for a multicentre, randomised, biomarker-stratified trial. BMJ Open 2018;8:e026767.

11. Kuwada T, Shiokawa M, Kodama Y, et al. Identification of an Anti-Integrin alphavbeta6 Autoantibody in Patients With Ulcerative Colitis. Gastroenterology 2021;160:2383–2394 e21.

12. Muramoto Y, Nihira H, Shiokawa M, et al. Anti-integrin alphavbeta6 antibody as a diagnostic marker for pediatric patients with ulcerative colitis. Gastroenterology 2022.

13. Rydell N, Ekoff H, Hellstrom PM, et al. Measurement of Serum IgG Anti-Integrin alphavbeta6 Autoantibodies Is a Promising Tool in the Diagnosis of Ulcerative Colitis. J Clin Med 2022;11.

14. Busk M, Pytela R, Sheppard D. Characterization of the integrin alpha v beta 6 as a fibronectin-binding protein. J Biol Chem 1992;267:5790–6.

15. Munger JS, Huang X, Kawakatsu H, et al. The integrin alpha v beta 6 binds and activates latent TGF beta 1: a mechanism for regulating pulmonary inflammation and fibrosis. Cell 1999;96:319–28.

16. Koivisto L, Bi J, Hakkinen L, et al. Integrin alphavbeta6: Structure, function and role in health and disease. Int J Biochem Cell Biol 2018;99:186–196.

17. Blanco-Mezquita JT, Hutcheon AE, Stepp MA, et al. alphaVbeta6 integrin promotes corneal wound healing. Invest Ophthalmol Vis Sci 2011;52:8505–13.

18. Huang XZ, Wu JF, Cass D, et al. Inactivation of the integrin beta 6 subunit gene reveals a role of epithelial integrins in regulating inflammation in the lung and skin. J Cell Biol 1996;133:921–8.

19. Yu Y, Chen S, Lu GF, et al. Alphavbeta6 is required in maintaining the intestinal epithelial barrier function. Cell Biol Int 2014;38:777–81.

20. Schmitz H, Barmeyer C, Fromm M, et al. Altered tight junction structure contributes to the impaired epithelial barrier function in ulcerative colitis. Gastroenterology 1999;116:301–9.

21. Porter CK, Riddle MS, Gutierrez RL, et al. Cohort profile of the PRoteomic Evaluation and Discovery in an IBD Cohort of Tri-service Subjects (PREDICTS) study: Rationale, organization, design, and baseline characteristics. Contemp Clin Trials Commun 2019;14:100345.

22. Silverberg MS, Satsangi J, Ahmad T, et al. Toward an integrated clinical, molecular and serological classification of inflammatory bowel disease: report of a Working Party of the 2005 Montreal World Congress of Gastroenterology. Can J Gastroenterol 2005;19 Suppl A:5A–36A.

23. Turpin W, Lee SH, Raygoza Garay JA, et al. Increased Intestinal Permeability Is Associated With Later Development of Crohn’s Disease. Gastroenterology 2020;159:2092–2100 e5.

24. Lennard-Jones JE, Shivananda S. Clinical uniformity of inflammatory bowel disease a presentation and during the first year of disease in the north and south of Europe. EC-IBD Study Group. Eur J Gastroenterol Hepatol 1997;9:353–9.

25. Shah S, Leleiko N, Lidofsky S, et al. Ocean State Crohn’s and Colitis Area Registry (OSCCAR): Incidence of Crohn’s Disease and Ulcerative Colitis in a Prospective, Population-Based Inception Cohort in Rhode Island: 1172. Official journal of the American College of Gastroenterology | ACG 2010;105:S425–S426.

26. Cohen BL, Zoega H, Shah SA, et al. Fatigue is highly associated with poor health-related quality of life, disability and depression in newly-diagnosed patients with inflammatory bowel disease, independent of disease activity. Aliment Pharmacol Ther 2014;39:811–22.

27. Therneau TM. A Package for Survival Analysis in R. https://CRAN.R-project.org/package=survival, 2020.

28. McCullagh P, Nelder JA. Generalized linear models. London ; New York: Chapman and Hall, 1989.

29. Robin X, Turck N, Hainard A, et al. pROC: an open-source package for R and S+ to analyze and compare ROC curves. BMC Bioinformatics 2011;12:77.

30. Dassopoulos T, Cohen RD, Scherl EJ, et al. Ulcerative Colitis Care Pathway. Gastroenterology 2015;149:238–45.

31. Duerr RH, Targan SR, Landers CJ, et al. Antineutrophil Cytoplasmic Antibodies in Ulcerative-Colitis - Comparison with Other Colitides Diarrheal Illnesses. Gastroenterology 1991;100:1590–1596.

32. Peeters M, Joossens S, Vermeire S, et al. Diagnostic value of anti-Saccharomyces cerevisiae and antineutrophil cytoplasmic autoantibodies in inflammatory bowel disease. American Journal of Gastroenterology 2001;96:730–734.

33. Israeli E, Grotto I, Gilburd B, et al. Anti-Saccharomyces cerevisiae and antineutrophil cytoplasmic antibodies as predictors of inflammatory bowel disease. Gut 2005;54:1232–6.

34. van Schaik FD, Oldenburg B, Hart AR, et al. Serological markers predict inflammatory bowel disease years before the diagnosis. Gut 2013;62:683–8.

35. Lochhead P, Khalili H, Ananthakrishnan AN, et al. Association Between Circulating Levels of C-Reactive Protein and Interleukin-6 and Risk of Inflammatory Bowel Disease. Clin Gastroenterol Hepatol 2016;14:818–824 e6.

36. Kevans D, Waterman M, Milgrom R, et al. Serological markers associated with disease behavior and response to anti-tumor necrosis factor therapy in ulcerative colitis. J Gastroenterol Hepatol 2015;30:64–70.

37. Hoie O, Aamodt G, Vermeire S, et al. Serological markers are associated with disease course in ulcerative colitis. A study in an unselected population-based cohort followed for 10 years. J Crohns Colitis 2008;2:114–22.

38. Kalla R, Adams AT, Bergemalm D, et al. Serum proteomic profiling at diagnosis predicts clinical course, and need for intensification of treatment in inflammatory bowel disease. J Crohns Colitis 2021;15:699–708.

39. Trzpis M, McLaughlin PM, de Leij LM, et al. Epithelial cell adhesion molecule: more than a carcinoma marker and adhesion molecule. Am J Pathol 2007;171:386–95.

40. de Lange KM, Moutsianas L, Lee JC, et al. Genome-wide association study implicates immune activation of multiple integrin genes in inflammatory bowel disease. Nat Genet 2017;49:256–261.

41. Uzzan M, Martin JC, Mesin L, et al. Ulcerative colitis is characterized by a plasmablast-skewed humoral response associated with disease activity. Nat Med 2022;28:766–779.

42. Breuss JM, Gallo J, DeLisser HM, et al. Expression of the beta 6 integrin subunit in development, neoplasia and tissue repair suggests a role in epithelial remodeling. J Cell Sci 1995;108 (Pt 6):2241–51.

43. Bandyopadhyay A, Raghavan S. Defining the role of integrin alphavbeta6 in cancer. Curr Drug Targets 2009;10:645–52.

44. Norris DA, Kissinger RM, Naughton GM, et al. Evidence for immunologic mechanisms in human vitiligo: patients’ sera induce damage to human melanocytes in vitro by complement-mediated damage and antibody-dependent cellular cytotoxicity. J Invest Dermatol 1988;90:783–9.

