## Supplementary Figure 1 and Tables for "Anti-integrin αvβ6 autoantibodies are a novel predictive biomarker in ulcerative colitis"

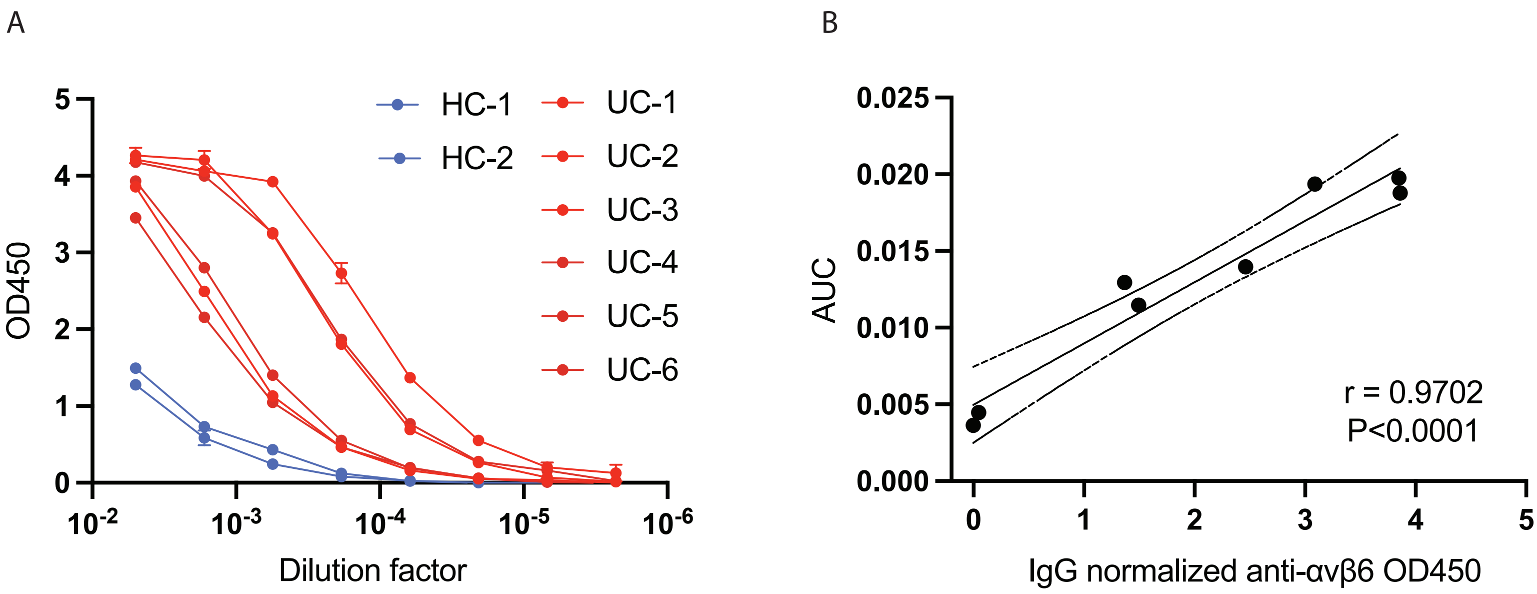


**Supplementary Figure 1: A. Anti-integrin αvβ6 IgG dilution curves in UC patients and normal volunteers. B. Pearson's correlation between IgG normalized OD450 anti-αvβ6 values and area under the curve (AUC) from dilution curves shown in A.**

**Supplementary Table 1: Conditional logistic regression models comparing anti-αvβ6 at each sample timepoint in those that developed UC compared to matched controls.**

| | OR | CI 95% (lower) | CI 95% (upper) | P value |
|---|---|---|---|---|
| Sample A anti-αvβ6 | 4.94 | 2.16 | 11.30 | 0.0002 |
| Sample B anti-αvβ6 | 24.88 | 4.22 | 146.75 | 0.0004 |
| Sample C anti-αvβ6 | 6.90 | 2.21 | 21.56 | 0.0009 |
| Sample D anti-αvβ6 | 23.99 | 3.37 | 170.82 | 0.0015 |

OR = Odds Ratio, CI = confidence interval

**Supplementary Table 2: Multivariate logistic regression model where ulcerative colitis status was modeled as function of anti-αvβ6 and multiple covariates such as age, gender and race based in the COMPASS cohort.**

| | OR | CI 95% (lower) | CI 95% (upper) | P value |
|---|---|---|---|---|
| Gender [ref: Male] Female | 1.04 | 0.15 | 7.02 | 0.97 |
| Age | 0.91 | 0.84 | 0.99 | 0.03 |
| Race [ref: White] Asian | 0.17 | 0.0005 | 57.19 | 0.55 |
| Race [ref: White] Black | 0.42 | 0.03 | 5.52 | 0.51 |
| Race [ref: White] Other | 0.50 | 0.06 | 4.21 | 0.52 |
| Anti-αvβ6 | 64.05 | 7.41 | 553.74 | 0.0002 |

OR = Odds Ratio, CI = confidence interval

**Supplementary Table 3: Multivariate logistic regression model where ulcerative colitis status was modeled as function of anti-αvβ6 and multiple covariates such as age, gender and race based in the OSCCAR cohort.**

| | OR | CI 95% (lower) | CI 95% (upper) | P value |
|---|---|---|---|---|
| Gender [ref: Male] Female | 2.93 | 0.80 | 10.70 | 0.10 |
| Age | 1.00 | 0.96 | 1.03 | 0.86 |
| Race [ref: White] Asian | 0.00 | 0.00 | Infinity | 0.99 |
| Race [ref: White] Black | 0.03 | 0.00 | 0.33 | 0.005 |
| Race [ref: White] Other | 0.02 | 0.00 | 0.43 | 0.01 |
| Anti-αvβ6 | 156.29 | 12.53 | 1949.04 | 0.0001 |

OR = Odds Ratio, CI = confidence interval

**Supplementary Table 4: Cox regression time to composite event with covariate of anti-αvβ6 in COMPASS cohort.**

| **Model 1** | | | | |
|---|---|---|---|---|
| Variable | HR | CI 95% (lower) | CI 95% (upper) | P value |
| anti-αvβ6 | 1.39 | 1.03 | 1.89 | 0.03 |
| Age | 0.99 | 0.96 | 1.02 | 0.37 |
| **Model 2** | | | | |
| anti-αvβ6 | 1.35 | 0.99 | 1.85 | 0.06 |
| Age | 0.99 | 0.96 | 1.02 | 0.46 |
| Extensive disease at baseline (E3) | 1.45 | 0.68 | 3.09 | 0.33 |

**Supplementary Table 5: Cox regression time to composite event with covariate of anti-αvβ6 in the OSCCAR cohort.**

| Variable | HR | CI 95% (lower) | CI 95% (upper) | P value |
|---|---|---|---|---|
| anti-αvβ6 | 1.24 | 1.01 | 1.53 | 0.04 |
| Age <40 years old at diagnosis | 1.05 | 0.57 | 1.95 | 0.87 |
| Extensive disease at baseline (E3) | 1.23 | 0.66 | 2.28 | 0.52 |
| ESR elevation at baseline | 0.70 | 0.32 | 1.53 | 0.38 |
| CRP elevation at baseline | 0.39 | 0.12 | 1.27 | 0.12 |
| Steroids at baseline | 1.64 | 0.84 | 3.23 | 0.15 |
| Focal ulcers on baseline endoscopy | 3.39 | 0.91 | 12.65 | 0.07 |
| Hospitalization at diagnosis | 1.27 | 0.59 | 2.76 | 0.54 |

**Supplementary Table 6: Cox regression time to composite event with covariate of pANCA in the OSCCAR cohort**

| Variable | HR | CI 95% (lower) | CI 95% (upper) | P value |
|---|---|---|---|---|
| anti-αvβ6 | 1.28 | 1.02 | 1.62 | 0.04 |
| pANCA | 0.75 | 0.36 | 1.54 | 0.43 |
| Age <40 years old at diagnosis | 1.10 | 0.59 | 2.05 | 0.76 |
| Extensive disease at baseline (E3) | 1.18 | 0.63 | 2.19 | 0.61 |
| ESR elevation at baseline | 0.69 | 0.32 | 1.50 | 0.35 |
| CRP elevation at baseline | 0.36 | 0.11 | 1.20 | 0.10 |
| Steroids at baseline | 1.69 | 0.86 | 3.32 | 0.13 |
| Focal ulcers on baseline endoscopy | 3.54 | 0.93 | 13.44 | 0.06 |
| Hospitalization at diagnosis | 1.21 | 0.55 | 2.63 | 0.64 |
